# Potential risk factors associated with human cystic echinococcosis: a semi-structured questionnaire from a large population-based ultrasound cross-sectional study in Eastern Europe and Turkey

**DOI:** 10.1101/575761

**Authors:** Francesca Tamarozzi, Okan Akhan, Carmen Michaela Cretu, Kamenna Vutova, Massimo Fabiani, Serra Orsten, Patrizio Pezzotti, Loredana Gabriela Popa, Valeri Velev, Mar Siles-Lucas, Enrico Brunetti, Adriano Casulli

## Abstract

**Background:** Cystic echinococcosis (CE) is a neglected parasitic zoonosis prioritized by the WHO for control. Hygiene education is included in CE control campaigns but appears of little impact, and the precise risk factors for human infection are still uncertain. Several works investigated potential risk factors for CE through questionnaires, mostly carried out on small samples, providing contrasting results. We present the analysis of risk factors questionnaires administered to participants to the largest prevalence study on CE conducted in Eastern Europe.

**Methodology/Principal Findings:** A semi-structured questionnaire was administered to 24,687 people from rural Bulgaria, Romania, and Turkey. CE cases were defined individuals with abdominal CE cysts detected on ultrasound. Variables associated with CE infection at p<0.20 in bivariate analysis were included into a multivariable logistic model, with a random effect to account for clustering at village level. Adjusted odds ratio (AOR) with 95%CI were used to describe the strength of associations. Data were weighted to reflect the relative distribution of the rural population in the study area by country, age group and sex. Valid records from 22,027 people were analyzed. According to the main occupation in the past 20 years, “housewife” (AOR 3.11 [1.51-6.41]) and “retired” (AOR 2.88 [1.09-7.65]) showed significantly higher odds of being infected compared to non-agricultural workers. “Having relatives with CE” (AOR 4.18 [1.77-9.88]) was also associated with higher odds of infection. Dog-related and food/water-related factors were not associated with infection.

**Conclusions/Significance:** Our results point to infection being acquired in a “domestic” rural environment and support the view that CE should be considered more a “soil-transmitted” than a “food-borne” infection, acquired through a “hand-to-mouth” mechanism. This result helps delineating the dynamics of infection transmission and have practical implications in the design of specific studies to shed light on actual sources of infection and inform control campaigns.

**AUTHOR SUMMARY:** Cystic echinococcosis (CE) is a parasitic disease with high socio-economic impact, mostly affecting pastoral communities. The causative agent, *Echinococcus granulosus sensu lato*, is naturally transmitted between dogs and livestock; humans acquire infection through accidental ingestion of parasite eggs. Hygiene education is among the strategies of CE control campaigns, but appears of little impact. “Ingestion of contaminated food/water”, and “contact with dogs” are generally mentioned as the sources of human infection, however actual risk factors are still undefined. Several works investigated potential risk factors for human CE infection through questionnaires, mostly carried out on small samples, providing contrasting results. We analysed 22,027 risk factors questionnaires administered to the participants of the largest prevalence study on CE conducted in Eastern Europe. We found that being “housewife” and “retired” as the main occupation in the past 20 years, and “having relatives with CE” were associated with higher odds of CE infection, while dog-related and food/water-related factors were not associated with infection. Our results indicate that CE may be considered more a “soil-transmitted” than a “food-borne” infection, acquired through a “hand-to-mouth” mechanism in a domestic, rural environment. This may help designing specific studies on pathways of transmission of this neglected parasite.

## INTRODUCTION

Cystic echinococcosis (CE) is a parasitic zoonotic disease caused by infection with the larval stage (metacestode) of the tapeworm *Echinococcus granulosus sensu lato* species complex. Its natural life cycle develops between canids (definitive hosts harbouring the adult stage in the intestine) and ungulates (intermediate hosts developing the larval stage in internal organs), in a predator-prey transmission pathway. The majority of human cases are documented in rural areas where livestock breeding is practiced, consistent with a life cycle mainly involving sheep and dogs [1, 2]. CE has remarkable health and socio-economic consequences for the rural populations affected [3, 4]. Current global estimates indicate a prevalence of 1-3 million cases of human CE, with a burden of 1-3.6 million Disability Adjusted Life Years and over 2 billion US$ costs accounting for human treatment and livestock production losses [4, 5]. In 2014, a joint FAO/WHO expert meeting ranked CE as the third most important food-borne parasitic disease at global level [6]. Further, in 2018 EFSA published a Scientific Opinion on public health risks associated with food-borne parasites, highlighting CE as “of the highest relevance in Europe” [7]. The WHO indicates CE as a zoonosis prioritized for control actions, including in Europe [8, 9].

Humans represent an accidental “dead-end” intermediate host for the metacestode of *E. granulosus*, thus they do not contribute to the perpetuation of the parasite’s life cycle. Only the implementation of preventive measures against infection, therefore, may reduce sustainably, in the long term, the burden of CE in humans in the presence of ongoing transmission in animals. This, in turn, may be controlled though implementation of abattoir surveillance and safe disposal of offal, culling of aged sheep, periodic deworming of dogs with praziquantel, and vaccination of sheep [10]. Hygiene education is one of the strategies included in CE control campaigns; however, on its own, this intervention did not impact significantly on transmission rate to humans, with the exception of Iceland [10].

Humans acquire infection through oral uptake of infective *E. granulosus* eggs; however, there is a great uncertainty on the actual source attribution and precise risk factors for infection. “Ingestion of contaminated food and water”, together with “direct contact/playing with dogs” are classically mentioned as the sources of human infection and are biologically plausible potential risk factors. However, actual data on contamination of and relative attribution from such sources are extremely scant and uncertain [7, 11, 12]. Further, Chaabane-Banaoues et al [13] found that degree of environmental contamination by *E. granulosus*-positive dog faeces did not necessarily correlate with human prevalence of CE, highlighting that multiple ecological factors, likely varying from area to area, and involving human behaviour and hygiene habits, are at the basis of human transmission.

Knowing the specific pathways of transmission to humans in endemic areas may allow optimizing and increasing the effectiveness of interventions aiming at the reduction of eggs ingestion by humans. However, the peculiarity of CE hamper this task. The absence of symptoms of “acute” human infection, the poorly known and apparently very long incubation time before the developing CE cyst(s) cause symptoms (if any), and the multiplicity of possible sources of infection, all contribute to the virtual impossibility to track back the infection episode(s) and specific associated risk behaviours.

Several works investigated the potential risk factors associated with human CE through questionnaires administered in hospital-based case-control and field-based cross-sectional studies, mostly providing contrasting results. Recently a systematic reviewed and meta-analysis by Possenti et al [14] aiming to summarize available data on statistically relevant potential risk factors, indicated that “living in endemic areas” and “dog ownership” seem to be the most significant potential risk factors for acquiring CE, consistently resulting from both case-control and cross-sectional studies. On the contrary, “dog contact” had a weak and non-significant association [14]. The same systematic review also found that factors related to habits involved in the perpetuation of the parasite life cycle (type of slaughtering, feeding dogs with raw viscera) were associated with increased risk of infection with variable statistical significance, while food- and water-borne pathways of transmission did not appear to impact significantly on human risk of acquiring CE [14]. Environmental contamination was identified as the main factor associated with CE infection also in more recent cross-sectional surveys carried out in endemic areas of Morocco [15] and Peru [16]. Factors associated with both parasite life cycle perpetuation and transmission through food or water were, on the contrary, reported as significantly associated with household risk of human CE in a recent Chinese study [17].

Previous questionnaire-based studies investigating potential risk factors associated with human CE generally tested samples of limited sizes and differed greatly one from the other for what concerns data collected in the interviews [14]. Furthermore, only about half of the field-based cross-sectional studies investigating risk factors through questionnaires used imaging as the diagnostic methodology for case definition, therefore confirming actual CE infection [14]. In 2014-2015, we carried out the largest research-based field survey on human CE in the context of the project “Human cystic Echinococcosis ReseArch in CentraL and Eastern Societies” (HERACLES [18]) funded by the European Commission in 2013. We examined by abdominal ultrasound 24,687 people from rural areas of Romania, Bulgaria, and Turkey, estimating that 151,000 people with abdominal CE may be currently present in the rural areas on these countries, a third of them potentially requiring treatment [19]. Here we present the analysis of risk factors questionnaires administered to participants during the HERACLES ultrasound population-based surveys.

## METHODS

### Ethics statement

Approval was granted by the Ethics Committees of Specialized Hospital for Active Treatment of Infectious and Parasitic Diseases “Prof. Ivan Kirov”, Sofia, Bulgaria; Colentina Teaching Hospital, Bucharest, Romania; and Hacettepe University Hospital, Ankara, Turkey.

### Ultrasound surveys

A detailed description of the US surveys has been published [19]. Briefly, abdominal ultrasound screening sessions were performed in 2014-2015 on 24,687 people in 50 villages of Bulgaria, Romania, and Turkey, in areas of mid-range endemicity for human CE. The screenings were carried out on the convenience sample of all volunteers living in the targeted endemic provinces who presented to the sessions in each village and signed the informed consent form. In all countries, a common protocol for diagnosis and clinical management of CE based on the WHO Informal Working Group on Echinococcosis (WHO-IWGE) Expert Consensus on clinical management of echinococcosis [20] was applied. Health education on CE was provided during the pre-screening project-advertising activities, and by the project team during the surveys with the aid of paper-based and audio-visual supports. Health education included information the life cycle of the parasite, ways of transmission to humans and key behaviours favouring the life cycle perpetuation, the characteristics of the disease in humans, and its treatment.

### Questionnaires

A semi-structured paper-based questionnaire (S1 Text) written in Bulgarian, Romanian, and Turkish was administered by the survey staff individually to each participant. Children, when appropriate, were helped answering by parents/guardians. The questionnaire was administered between the signature of the informed consent form and the ultrasound exam. The questionnaire included demographic, occupational, and schooling-related questions, as well as questions concerning knowledge about existence of human CE and occurrence of cases in the family, dog and livestock-related practices, and food- and drinking water-related habits. After the field surveys, data were transferred to an electronic database (Excel), and manually curated before analysis. The complete list of variables for analysis is presented in S1 Table. The analysis of the answers to the question “years of school attended” was carried out in four “Education” categories. Similarly, the question relative to dog’s treatment with praziquantel was analysed in four categories. Occupations were grouped into six categories. The answers to the questions “Do you leave dogs free to roam” and “What do you feed dogs with” were double-checked for incongruences and manually curated based on answers to the related questions “Reasons for keeping dogs” and “How do you dispose of viscera”. Similarly, the answers whether the interviewed carried out agricultural activities in the past 20 years were double-checked for incongruence and manually curated based on current and past occupation. Ways of disposal of viscera from slaughtered animals was analysed irrespective of whether the person carried out home slaughter, as it is common practice in the investigated areas to obtain livestock viscera that are fed to dogs even without owning livestock and/or slaughtering livestock at home. Finally, due to the extreme heterogeneity of the answers provided, we could not analyse the answers to the questions addressing the frequency of antiparasitic treatment of dogs, the recognition of the picture of a CE cyst in the liver of a sheep, the time of the year in which home slaughter was carried-out, the ways of disposal of dogs’ feces, and the access of dogs not owned by the interviewed in his/her household facilities.

### Case definition

A detailed description of classification of patients and CE cysts has been published [19]. In brief, infection with CE was based on ultrasound imaging, evaluated by two sonographers during the screening and confirmed by re-evaluation of each lesion through images and video files before data analysis. Cysts were identified and staged based on the visualization of pathognomonic signs of CE etiology, according to the WHO-IWGE Expert Consensus [20]; more stringent conditions were, however, applied to unilocular cysts, which where ascribed to parasitic etiology only if a double wall was clearly visible. Lesions suspect of CE, including CL, were investigated as per protocol to define the nature of the lesion [19]. Due to logistic constraints, only patients visiting the project’s referral hospitals for treatment of CE cysts received chest X-ray for the detection of possible lung CE; none resulted positive [19]. “CE cases” were defined as all individuals with abdominal CE cysts detected on ultrasound, independently of whether they reported having received previous treatment for CE.

### Statistical analysis

We excluded from the analysis questionnaires from 128 (0.5%) individuals who had no CE cysts detected by imaging, but self-reported treatment for CE in the past, as it was not possible to confirm their infection through clinical documentation. Moreover, we excluded questionnaires with incomplete information (n=2426; 9.8%) or unresolvable incongruences n=(106; 0.4%), leaving complete records from 22,027 (89.3%) participants available for the analysis. Individuals (n=38) with suspect lesions, the etiology of which could not be ascertained, were considered as CE-negative. We described the socio-demographic characteristics and risk profile of the study sample population through counts and percentages. The prevalence of CE was estimated using sampling weights to reflect the relative distribution of the rural population in the study area by country, age group and sex, as derived from official population statistics [19]. Prevalence estimates were presented with 95% confidence intervals (CI), calculated using the Taylor linearization method to account for the increased variance due to the sampling design. The association between the presence of CE infection and each potential risk factor was evaluated using the χ2 test on the whole 22,027 questionnaires sample, based on the geographical and ecological contiguity of the entire investigated area. All variables associated with CE infection at p<0.20 in bivariate analysis were included into a multivariable logistic model together with a random effect to account for clustering at village level. Regardless of its association with CE infection in bivariate analysis, we excluded “current occupation” from the multivariable analysis to prevent collinearity problems due to its strong association with the variable “prevalent occupation in the past 20 years” (we assumed the latter as more appropriate to evaluate the occupation-related risk for an infection that was likely acquired years previously). We scaled sampling weights according to the actual clusters’ size before running the multilevel model [21]. The adjusted odds ratio (AOR) with 95% CI were used to describe the strength of the associations. The Interactions between each risk factor and country were assessed through the Wald test. Country-specific multivariable models were also computed. Finally, we used the intraclass correlation coefficient (ICC) to estimate the proportion of the residual variability attributable to the village-related context [22]. Statistical significance was set at p<0.05. The analysis was performed using Stata/MP version 14.2 (StataCorp LP, College Station, Texas, USA).

## RESULTS

Of 22,027 analysed questionnaires, 13,957 (63.4%) belonged to females and 8070 (36.6%) to males; 105 (0.51%, 95% CI: 0.24-1.07) people had abdominal CE during the ultrasound screenings (n=71 [0.63%, 95% CI: 0.25-1.55] females and n=34 [0.39%, 95% CI: 0.23-0.67] males). The results of the descriptive and bivariate analysis performed on the whole sample are presented in S1 Table. Ten of the 23 analysed variables were associated with CE infection at p<0.20 and included into the multivariable logistic model.

The results of the multivariable analysis are presented in Table 1 and graphically depicted in Figure 1. In relation to the main occupation in the past 20 years, housewives (AOR 3.11; 95% CI 1.51-6.41; p=0.002) and retired persons (AOR 2.88; 95% CI 1.09-7.65; p=0.033) showed an increased odds of infection compared to non-agricultural or office/service workers. Having had relatives with CE was positively associated with CE infection (AOR 4.18; 95% CI 1.77-9.88; p=0.001), while individuals with university or higher level of education showed a significantly reduced odds of infection compared to those without any formal education (AOR 0.11; 95% CI 0.01-0.88; p=0.038). Other factors were associated with an increased odds of CE infection but results were only borderline statistically significant (p<0.1). These were: “Farmer\livestock breeder\other agricultural or veterinary activities as the main occupation in the past 20 years” (AOR 2.49; 95% CI 0.93-6.66; p=0.068) and “Giving raw viscera to dogs” (AOR 1.50: 95% CI 0.95-2.38; p=0.080). “Drinking commercial water” was associated with a reduced odds of CE infection with borderline significance (AOR 0.65; 95% CI 0.40-1.04; p=0.071).

**Table 1.** Results of the multilevel logistic regression model, including village as random effect. *Accounting for clustering at village level. **Adjusted OR per linear 10-years increase in age. #Statistically significant interaction with country (Wald test p<0.05)

| Variable | Adjusted OR | 95% CI* | p-value* |
| --- | --- | --- | --- |
| SEX |  |  |  |
| Female | 1 |  |  |
| Male | 0.98 | 0.64-1.5 | 0.930 |
| AGE** | 1.05 | 0.89-1.23 | 0.582 |
| LIVED IN AREAS WITH HIGH DENSITY OF DOGS AND SHEEP IN THE PAST 20 YEARS |  |  |  |
| No | 1 |  |  |
| Yes | 1.94 | 0.84-4.47 | 0.118 |
| MAIN OCCUPATION IN THE PAST 20 YEARS# |  |  |  |
| Non-agricultural activities or office/service employee | 1 |  |  |
| **Housewife** | **3.11** | **1.51-6.41** | **0.002** |
| Farmer/livestock breeder/other agricultural/veterinary activities | 2.49 | 0.93-6.66 | 0.068 |
| Students and children <5 years of age | 1.31 | 0.61-2.85 | 0.487 |
| **Retired** | **2.88** | **1.09-7.65** | **0.033** |
| Unemployed | 2.09 | 0.51-8.58 | 0.309 |
| AGRICULTURAL ACTIVITIES IN THE PAST 20 YEARS |  |  |  |
| No | 1 |  |  |
| Yes | 1.47 | 0.64-3.36 | 0.366 |
| EDUCATION |  |  |  |
| None | 1 |  |  |
| Primary | 1.05 | 0.55-2.00 | 0.893 |
| Secondary/High school | 1.15 | 0.60-2.18 | 0.678 |
| **University/Postgraduate** | **0.11** | **0.01-0.88** | **0.038** |
| KNOWLEDGE OF HUMAN CE EXISTENCE |  |  |  |
| No | 1 |  |  |
| Yes | 1.78 | 0.69-4.58 | 0.235 |
| KNOWN PRESENCE OF RELATIVES WITH CE |  |  |  |
| No | 1 |  |  |
| **Yes** | **4.18** | **1.77-9.88** | **0.001** |
| RAW VISCERA GIVEN TO DOGS |  |  |  |
| No | 1 |  |  |
| Yes | 1.50 | 0.95-2.38 | 0.080 |
| DRINK COMMERCIAL WATER<sup>#</sup> |  |  |  |
| No | 1 |  |  |
| Yes | 0.65 | 0.40-1.04 | 0.071 |

**Fig 1.**
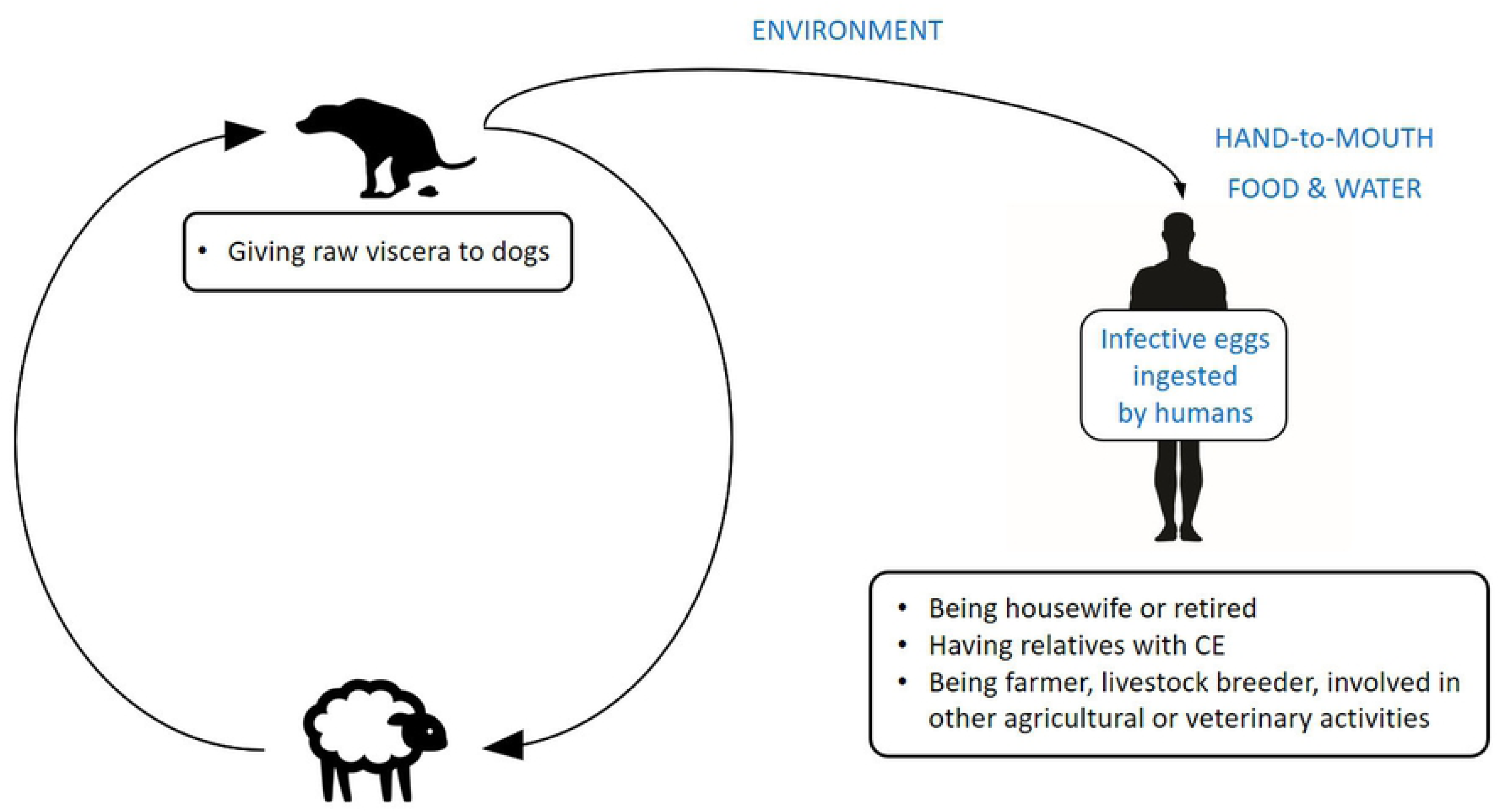
Schematic representation of *E. granulosus* life cycle, pathways of transmission to humans (in blue), and potential risk factors associated with increased odds of human CE infection identified in our study (bullet points).

The estimate of the intraclass correlation coefficient (ICC=16.6%; 95% CI 7.2-33.7) indicates that almost one fifth of the residual variability not explained by the individual-level variables included into the multivariable model was likely due to village-related contextual factors.

A significant interaction with country was found only for “Main occupation in the past 20 years” and “Drinking commercial water” (Wald test, p<0.05). The results of the country-specific multivariable models are presented in S2 Table. In general, these were consistent with results from the analysis of the whole sample. However, in Bulgaria, a statistically significant increase in odds of CE infection was observed for “Student or children <5 years as the main occupation in the past 20 years” (AOR 3.04; 95% CI 1.10-8.32; p=0.032) and individuals with “primary-level” highest education (AOR 2.98; 95% CI 1.15-7.69; p=0.024), while these associations, although not statistically significant, appeared reversed in Romania and Turkey. Moreover, a significantly increased risk of being infected was observed in individuals reporting past agricultural activities in Turkey (AOR 2.91; 95% CI 1.50-5.67; p=0.002), but not in those living in Romania and Bulgaria. All the other investigated associations were found to be not statistically significant or having the same direction in all countries.

## DISCUSSION

WHO advocates control of CE [8]. Reference control strategies include “health education”; however, the content and target population of such educational intervention(s) varied between campaigns and, overall, did not appear having significantly affected the transmission of CE to humans [10]. Knowing more precisely human infection risk factors in endemic areas may allow optimizing and modelling hygiene-based educational interventions aiming at the reduction of eggs ingestion by humans. However, this is particularly difficult due to the absence of symptoms of “acute” human infection and the unknown, likely months to years-long, interval between infection and diagnosis. Multiple potential habits/sources may result in human ingestion of infective parasite eggs. However, so far very few experimental data are available on the actual contamination of different materials by *E. granulosus* eggs [12], and the analyses of questionnaires investigating potential risk factors gave contrasting results [14]. Our questionnaire-based study, carried out in the context of the largest research-based cross-sectional survey on human CE [19], applying stringent case definition, may help better framing the general characteristics of risk factors for human infection.

Owned dog-related factors (owing dogs and length of dog ownership, reason for keeping dogs, allowing dogs to roam or enter the house, antiparasitic treatment of dogs) were not found associated with odds of human infection. This may result from the dog husbandry habits in the investigated areas, which may allow environmental contamination with parasite eggs even outside the premises of the interviewed person; also a variable meaning of “owing” a dog in different areas, as also noted in previous studies, may have influenced this result [15]. Food/drinking water-related factors such as consumption of unwashed vegetables and of potentially unsafe water were also not associated with increased odds of human infection. While these results indicate that food- and water-borne transmission may not play a major role, increased risk of infection deriving from occupations (housewife, retired) related to the household point to infection being acquired in a “domestic” rural environment. This is also supported by the increased odds of infection associated with having an agricultural-related occupation. Increased risk of infection associated with having relatives with CE, as well as the trend toward in increased risk associated with having some knowledge of the existence of human CE, may derive from living in a context where human CE and therefore where its transmission cycle is common. This transmission cycle is perpetuated by the habit of giving raw viscera to dogs, individuated as a risk factor of borderline significance in our analysis, as it induces dog infection and in turn environmental contamination through shedding of infected feces. The fact that another habit potentially favouring the transmission of *E. granulosus* to dogs, that is home slaughtering of livestock, was not associated with increased odds of infection may be due to the common habit of obtaining viscera to feed dogs even if the household did not own and/or did not carry out informal slaughter, as also occurs in other geographical areas [15]. The risk deriving from environmental contamination appears reduced by socio-economic related factors: in our analysis, having a high education and drinking commercial water were associated with reduced odds of infection. Lack of association with sex and age could be explained by the common exposure, in rural areas, of both sexes and at all ages. However, although not statistically significant, an increase in infection prevalence with age can be anyway observed in our data set, as expected for a chronic infection.

Our results are overall in line with those of the systematic review of Possenti and colleagues [14], and with the recent work carried out in Peru and Morocco [15, 16], individuating environmental contamination and likely “hand-to-mouth” transmission as the main factor responsible for CE transmission, while food/water source attribution likely not of primary importance.

Our study has some limitations deserving discussion. First, limitation deriving from recall bias is intrinsic to the study design and the peculiarity of a chronic, often asymptomatic, infection such as CE. Second, it is possible the health education information provided before the administration of the questionnaire would have influenced the answer to the question related to knowledge of CE in humans. However, if this was systematically the case, it would have rather resulted in no association between infection and answer to this question, contrary to what we found. Third, it was difficult to ascertain whether participants replied to questions in terms of their “common habits” or “even occasional” behaviours. This was evident from the mismatch observed between related questions such as: “Do you leave dogs free to roam – reply “No”” and “Reasons to keep dogs – reply “hunting” or “herding””; “What do you feed dogs with – reply “only commercial/cooked food”” and “How do you dispose of viscera – reply “give raw to dogs””; and “Agricultural activities carried out in the past 20 years – reply “No” and agricultural-related current and/or past occupations. These problem could have been reduced by pre-testing the questionnaire, which was not carried out due to time constraint deriving from the organization timeline of ultrasound surveys. Another limitation may derive from the possible inclusion of some CE-infected individuals among the non-infected group. This may have derived from the stringent case definition applied in the survey [19], and the impossibility of performing a chest radiograph to all survey participants for the detection of isolated lung cyst. Finally, it is worth highlighting that the variables included in our questionnaires were not investigating infection transmission behaviours directly, but can be regarded as indirect driving factors related to socio-economic status and hygiene-related habits.

To conclude, our study carried out in the context of the largest research-based cross-sectional study conducted on CE [19], contributes to the scientific knowledge of potential risk factors of human CE. Our results support the paradigm shift view that CE should be mainly considered an “environmental-borne” infection, similar to the “classical” soil-transmitted helminthiases, transmitted through a “hand-to-mouth” mechanism, while food/water-borne transmission may be of secondary importance. In both cases, however, the “community risk” in endemic areas should be highlighted, aside of “individual” risk factors. These concepts, supported by our results, are pivotal in delineating the general dynamics of infection transmission and have important practical implications for public health policy makers across endemic countries in the design control campaigns. However, more country/community-specific and habits-specific questionnaires, as well as experimental studies on parasite contamination of matrices, are needed to shed light on actual sources of infecting eggs and on behaviours at risk for individual infection.

## ACKNOWLEGDMENTS

We are grateful to Prof Thomas Junghanss (Head of the Section of Clinical Tropical Medicine at Heidelberg University Hospital and Chair of the WHO Informal Working Group on Echinococcosis), Bernadette Abela-Ridder (Team Leader, Neglected Zoonotic Diseases, Department of the Control of Neglected Tropical Diseases, WHO), and Johanna Takkinen (Head of Disease Programme Food- and Waterborne Diseases and Zoonoses, European Centre for Disease Prevention and Control) for supporting this research. We are thankful to the staff who participated to the field work in Bulgaria (Rossitza Chipeva, Branimir Golemanov, Marin Muhtarov), in Romania (Corina Manuela Constantin, Denisa Janta, Patricia Mihailescu, Marius Petrutescu, Alexandru Cosmin Popa, Mircea Ioan Popa), and in Turkey (Devrim Akinci, Turkmen Ciftci).

We also thank Madalina Preda, Adelina Silvana Gheorghe, Andrei Capraru, Luca Stan Sion and Zlatka Todorova Muhtarova, who volunteered to help during the fieldwork.

## SUPPORTING INFORMATION LEGENDS

**S1 Text.** Semi-structured paper-based questionnaire (English version).

**S1 Table.** Results of the bivariate analysis performed on the whole sample. *Calculated by weighting data according to the rural population size by country, sex and age group. ** Accounting for clustering at village level. NC, not calculable.

**S2 Table**. Results of the multilevel logistic regression model, including village as random effect. *Accounting for clustering at village level. **Adjusted OR per linear 10-years increase in age. °BULGARIA: variance between villages 0.21 (95% CI 0.02-2.17), intra-village correlation 6.0 (95% CI 0.6-39.7); ROMANIA: variance between villages 0.30 (95% CI 0.03-3.51), intra-village correlation 8.3 (95% CI 0.8-51.6); TURKEY: variance between villages 1.48 (95% CI 0.32-6.96), intra-village correlation 31.1 (95% CI 8.8-67.9).

**S1 Checklist**. STROBE checklist

